# Preclinical Evaluation of PSMA-Targeted Ultrasound Contrast Agents in an Orthotopic Model of Prostate Cancer in Rabbits

**DOI:** 10.1101/2025.08.07.668901

**Authors:** Felipe M. Berg, Eric C. Abenojar, Pinunta Nittayacharn, Nathan K. Hoggard, Sidhartha Tavri, Jing Wang, Felipe Matsunaga, Michaela B. Cooley, Dana Wegierak, Xinning Wang, Thomas J. Rosol, James P. Basilion, Agata A. Exner

**Author notes:** **Correspondence to:** Prof. Agata A. Exner, PhD, Department of Radiology and Department of Biomedical Engineering, Case Western Reserve University, 10900 Euclid Avenue, Cleveland, OH, 44106, USA. These authors contributed equally to this work (Felipe M. Berg, Eric C. Abenojar).

## Abstract

The localization of prostate cancer by ultrasound remains limited by the lack of B-mode conspicuity and the confinement of clinically approved microbubbles (MBs) to the vasculature. This precludes the differentiation of viable tumor tissue, necrotic tissue, and margin-associated disease. We investigate the use of prostate-specific membrane antigen (PSMA)-targeted lipid-shelled perfluorocarbon nanobubbles (PSMA-NBs) in an orthotopic rabbit model using a clinical CEUS system. We implanted PSMA-positive PC3pip-GFP tumors into the prostates of immunosuppressed New Zealand White rabbits and performed transabdominal imaging with PSMA-NBs, MBs, and untargeted nanobubbles (Plain-NB) using the same imaging and analysis protocols. To address tumor heterogeneity and ultrasound boundary ambiguity, ROIs were defined from baseline B-mode images and segmented into the tumor core, rim, and a surrounding peritumoral area. Pixel-wise parametric and decorrelation time (DT) maps were generated and correlated with whole-slide histology (H&E) and PSMA IHC. Compared to MBs, PSMA-NBs exhibited higher peak intensities in the tumor core and rim (1.60-fold and 1.50-fold, respectively) and improved retention (mean transit time: 4.20 to 4.50-fold higher) for up to 10 minutes in the tumor and peritumoral areas. PSMA-NB kinetic and DT parameters also correlated with histology-defined tumor viability and PSMA expression. Compared to Plain-NBs, PSMA-NBs also exhibited improved retention (MTT +21%; AUCwo) in the rim and peritumoral areas. This study demonstrates the capability of PSMA-NBs to characterize prostate cancer by molecular CEUS beyond what is possible with conventional MBs.

## Introduction

Prostate cancer is the second leading cause of cancer-related deaths in men in the United States, with over 30,000 deaths occurring every year.^1^ Despite the progress made in the diagnostic and therapeutic modalities, the financial burden of prostate cancer exceeds $22.3 billion, making it imperative to develop more accurate and cost-effective diagnostic modalities.^2^ The conventional diagnostic modalities involve the use of ultrasound-guided prostate biopsy in patients with elevated levels of prostate-specific antigen or abnormal digital rectal examination findings.^3–5^ Magnetic resonance imaging (MRI)-US fusion-guided biopsies are considered the gold standard, but their limited availability outside of Europe and North America means that B-mode US remains the primary worldwide method for guiding prostate biopsies.^4–7^ However, the low sensitivity and resolution of the ultrasound modality to distinguish between normal and malignant tissues make it necessary to obtain multiple biopsy cores, thereby increasing the risk of complications, including significant infection rates.^8–10^

To improve soft tissue contrast and structure visualization with US, microbubble (MBs) ultrasound contrast agents (UCA) were introduced, primarily for cardiovascular applications.^11,12^ More recently, FDA-approved UCAs (e.g. Lumason^®^) have been employed for the detection of neoplastic lesions.^13^ However, the intravascular nature of MBs, due to their 1-5 μm diameter, restricts their utility in tumors to the vasculature. However, there may be a benefit to additional assessment beyond vascular characteristics. Nanobubble (NB) UCAs were developed as an alternative to MBs and can provide complementary information that has the potential to probe the extravascular tumor biology.^14,15^ With diameters of 200-500 nm, NBs can extravasate of pathologically permeable blood vessels, commonly seen in inflammatory conditions, providing information about the surrounding environment and parenchyma.^16–18^

As tumors grow, they frequently develop a necrotic core due to inadequate vascular supply.^19,20^ This core, characterized by metabolic stress and inflammation, is a hallmark of advanced disease and presents a significant challenge for imaging technologies, which struggle to delineate tumor boundaries and differentiate between viable and necrotic tissue.^19,21^ An ideal contrast agent should not only identify the tumor, but also distinguish its regions, including the core, rim, and surrounding inflamed tissue, to improve both diagnostic accuracy and therapeutic planning. NBs are excellent candidates for this purpose due to their ability to extravasate and linger in the tissues.^16,22^

Prostate-specific membrane antigen (PSMA) is a molecular target in prostate cancer that has been validated in the clinic by its high-level expression in a significant proportion of clinically significant prostate cancer and metastatic sites, as well as its established role in imaging and therapeutic interventions.^23,24^ In current clinical practice, PSMA PET has become a mainstay in the evaluation of prostate cancer patients, including the initial evaluation of those with unfavorable intermediate- and high-risk prostate cancer, as well as the evaluation of patients with suspected biochemical recurrence.^24^ In these contexts, the modality is often found to impact the overall management strategy by increasing the sensitivity to detect nodal and metastatic sites compared to other imaging modalities.^24^ Moreover, PSMA-targeted theranostics, such as radioligand therapy using PSMA-targeting agents, underscore the importance of PSMA expression and the need to accurately characterize PSMA in prostate cancer.^23^

In the context of the current research, PSMA-targeted NBs (PSMA-NBs), which have shown promising results in preclinical studies,^25–27^ could potentially further improve the accuracy of prostate cancer imaging before, during and after an interventional procedure such as prostate biopsy or focal therapy. The small hydrodynamic diameter of these nanobubbles enables their extravasation into the extracellular matrix and their retention in PSMA-expressing tissues for long periods compared to MBs, especially in the setting of molecular targeting.^26^ This is particularly relevant because ultrasound remains broadly accessible and real-time, yet lacks an established clinically deployed molecular imaging analogue to PSMA PET; a PSMA-targeted CEUS approach could therefore complement current imaging workflows by enabling bedside, real-time assessment of PSMA-expressing disease. The long retention times are essential because they create opportunities for both imaging and therapeutic applications of these NBs. Previous studies have shown that prostate cancer cells expressing PSMA can be selectively visualized using PSMA-NBs in the setting of ultrasound molecular imaging, which is mediated by pharmacokinetics that are distinct from those of plain/untargeted NBs. These NBs interact with cell surface receptors and are internalized via receptor-mediated endocytosis.^26–28^ However, the application of these results is limited by the use of mouse models in these studies,^25,28–30^ which do not adequately mimic the prostate cancer environment in humans.^31^

The present study aims to validate PSMA-NB CEUS in a large animal model that is more comparable to human prostate cancer, and can accommodate an orthotopic tumor to enable the comparison of agent activity and detection sensitivity in cancerous versus normal prostate tissue. While orthotopic mouse models have been explored for this purpose, the murine normal prostate has a highly distinct anatomy and it cannot accommodate the size of lesions which would be of clinical relevance, and the tumors do not develop in a manner that is consistent with human prostate cancers. Following a previously established rabbit orthotopic model,^32^ here we aim to demonstrate the ability of PSMA-NBs to differentiate between necrotic and vital tumor areas, PSMA-expression in tumors as well as normal prostate tissue, improving diagnostic accuracy. With the validation of PSMA-NBs in a large animal model, we are a step closer to clinical trials.

## Methods

**Industry Support:** Siemens^®^ Healthineers (Erlangen, Germany) provided the ultrasound system used in this study (Acuson S3000 with 18L6 transducer). The company had no involvement in study design, data analysis, or manuscript preparation. All authors had full control of the data and submission, and none are affiliated with Siemens^®^ Healthineers.

**Ethics Approval:** All animal procedures were approved by the Institutional Animal Care and Use Committee (IACUC) and were conducted in accordance with the National Institute of Health (NIH) guidelines and relevant institutional regulations for the care and use of laboratory animals.

### Ultrasound Contrast Agent Preparation

The preparation and characterization of the Plain-NB (untargeted) and PSMA-NBs (targeted) contrast agents followed established protocols.^14,26^ In summary, the lipids DBPC (Avanti Polar Lipids Inc., Pelham, AL, United States), DPPE (Corden Pharma, Switzerland), DPPA (Corden Pharma, Switzerland), and mPEG-DSPE2000 (Laysan Lipids, Arab, AL, United States) were dissolved in a mixture with propylene glycol (PG, Sigma Aldrich, Milwaukee, WI, United States), and glycerol (Thermo Fisher Scientific, Fair Lawn, NJ, United States). Following dissolution, the lipids were dispersed in prewarmed phosphate-buffered saline (PBS, Gibco Life Technologies, Grand Island, NY, United States) to form an oil-in-water emulsion. The emulsion was placed in a 3 mL vial and sealed, and the vial was evacuated and filled with perfluoropropane (C_3_F_8_, AirGas, Radnor, PA, United States) gas. For PSMA-NB, 25 μL of DSPE-PSMA-1 at a concentration of 1 mg/mL was added into the vial along with the lipid dispersion prior to evacuation. Following the gas exchange process, the vial was agitated using a Vialmix^®^ device (Lantheus Medical Imaging, Canada) for 45 seconds, inverted, and centrifuged at 50 RCF for 5 minutes. 400 microliters of the resultant Plain-NB or PSMA-NBs were collected from each vial, and multiple vials were combined to achieve the required volume for imaging studies. A flowchart summarizing the NB synthesis process is provided in Supplementary Figure S1. Characterization using resonant mass measurement (RMM) and dynamic light scattering (DLS) demonstrated that PSMA-NBs have a hydrodynamic diameter of 266–288 nm, a concentration of 3.9 x 10^11^ ± 2.82 x 10^10^ NBs/ml, and a slightly negative zeta potential, which ensures stability. For comparison, untargeted NBs were shown to have an average diameter of 274 ± 8 nm with a concentration of 4.07 x 10^11^ ± 3.15 × 10^10^ NBs/mL. Furthermore, *in vitro* cellular uptake experiments defined the optimal PSMA ligand density as incorporation of 25 μg of PSMA-1 into the formulation, corresponding to approximately 35 x 10³ PSMA molecules per NB and yielding robust receptor-mediated targeting.^14,28^ For comparative purposes, commercially available Lumason^®^ MBs (Bracco, Italy) were prepared according to the manufacturer’s instructions.^33^

### Cell Culture

PSMA-positive human PC3pip cells (human prostate cancer) were retrovirally transduced^34^ and subsequently modified to express green fluorescent protein (GFP). The authenticity of the cell line was previously verified via Western blot analysis.^35^ These cells were maintained in a humidified incubator at 37°C and 5% CO_2_ in RPMI 1640 medium (Gibco Life Technologies, Grand Island, NY), supplemented with 10% fetal bovine serum (FBS - Invitrogen Life Technology, Grand Island, NY) and 1% penicillin/streptomycin (Gibco Life Technologies, Grand Island, NY).

### Tumor Inoculation

All animal procedures were performed according to protocols approved by the Institutional Animal Care and Use Committee (IACUC), adhering to relevant guidelines and regulations. Immunosuppression was induced with daily subcutaneous injections of cyclosporine (10 mg/kg), starting one day prior to tumor cell injection and continued until the study concluded (Figure 1A). Nine sexually mature male New Zealand White rabbits (Charles River Laboratories, Ashland, OH, United States) were utilized. Anesthesia was induced and maintained with 1–2% isoflurane throughout all procedures and vital signs were monitored for procedure duration and recovery from anesthesia. For tumor inoculation, rabbits were placed in a supine position, and the lower abdomen was shaved and prepared using aseptic techniques. After shaving, additional hair was removed with potassium thioglycolate cream (Veet Industries, Warren, MI). The prostate gland was located using US, and 8 x 10^6^ PC3pip GFP cells suspended in PBS were injected into the prostate gland using a 50.8 mm, 21-gauge PrecisionGlide™ needle (BD, Franklin Lakes, NJ, United States). The needle was introduced percutaneously and advanced to the prostate while avoiding the iliac vessels using US guidance by an interventional radiologist (Figure 1B). Once the needle tip reached the central portion of the prostate, 100 µL of the cell suspension was injected. Both the syringe and needle were primed with cells prior to injection.^32^

**Figure 1.**
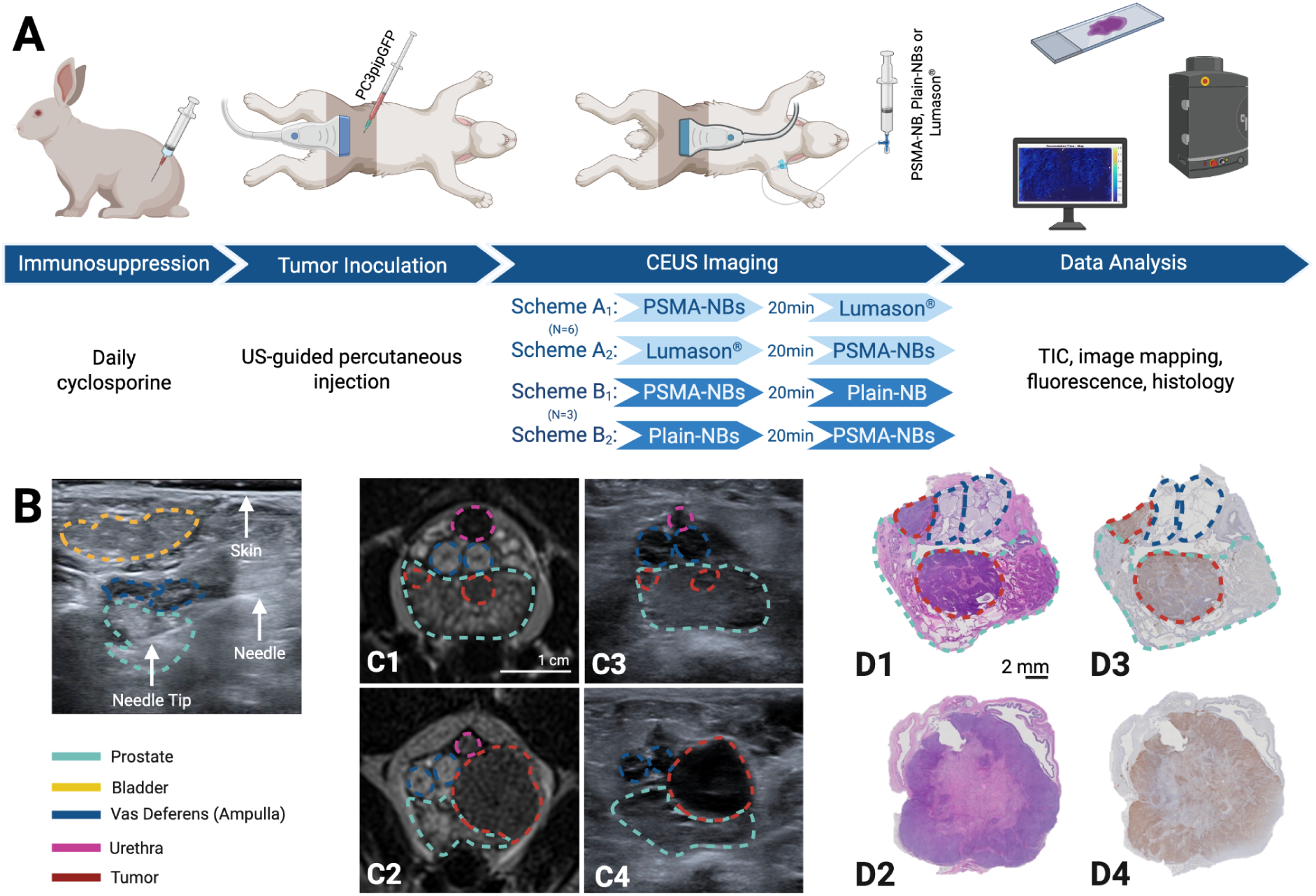
Study protocol summary. (A) Schematic diagram of the experimental setup and study methodology. (B) Representative image of the percutaneous tumor cells inoculation and relevant anatomical structures. (C1-2) Representative images of magnetic resonance imaging, (C3-4) B-mode ultrasound, (D1-2) H&E histology and (D3-4) prostate-specific membrane antigen (PSMA) immunohistochemistry (IHC) for intraprostatic tumors (C1, C3, D1, D3) and extraprostatic tumors (C2, C4, D2, D4).

### B-Model Ultrasound and MRI

Ultrasound imaging, including B-mode and Doppler, for guided tumor cell injection and subsequent weekly monitoring, was conducted using an Acuson S3000 US scanner (Siemens Healthineers, Germany) equipped with an 18 MHz (18L6) transducer. Trans-abdominal images were acquired. MRI scans were performed using a 3T Magnetom Vida scanner (Siemens Healthineers, Germany) with the following parameters: echo time (TE) = 85 ms, repetition time (TR) = 7000 ms, flip angle = 160°, bandwidth = 203 Hz/pixel, voxel size = 0.4 x 0.4 x 2.0 mm^3^, field of view (FOV) = 120 x 86.3 mm (sagittal) and 120 x 97.5 mm (transverse), base resolution = 320, 26 slices. A board certified abdominal radiologist interpreted the MRI images to determine the tumor’s location and size. Rabbits were anesthetized with 2% isoflurane during US procedures. For the MRI, rabbits were induced with 2% isoflurane and anesthetized with ketamine (10 mg/kg) intramuscularly 10 minutes before scanning. Additional ketamine (5 mg/kg) was injected intravenously under veterinary supervision as needed to maintain sedation.

### Contrast Enhanced Ultrasound Imaging

Contrast-enhanced ultrasound (CEUS) imaging, using the same US system described above, was performed once a week and employed the standard nonlinear contrast pulse sequencing (CPS) mode with the following settings: frequency 8.0 MHz, mechanical index (MI): 0.13, dynamic range (DR): 55 dB, gain: −5 dB, and 1 frame per second (fps). Plain-NB, PSMA-NBs and MBs at a dose of 0.8 mL/kg were injected intravenously via an ear vein catheter. CEUS imaging was conducted continuously for 10 minutes, with a 15-second background recording prior to bubble injection (Video 1). Rabbits were alternately injected with PSMA-NBs and MBs (N=6) or Plain NB (N=3) each week (Figure 1A), with the order of injection reversed every week. A 20-minute clearance period was maintained between the administration of each UCA (Figure 1A).

### Animal Enrollment, Exclusion Criteria, and Data Acquisition

One animal was euthanized at 2 weeks due to gastrointestinal complications related to cyclosporine, in compliance with IACUC and the Animal Welfare Act guidelines and therefore not included in the analysis. 14 tumors were successfully induced in the eight remaining enrolled animals. At each weekly session, we acquired contrast data from only one 2D plane. As a result, animals with multiple spatially separated tumors could not have all lesions imaged at every time point.. In some scans, the prostate had to be excluded from the imaging plane to capture the tumor, resulting in scans that contained only tumor images. In total, 14 tumor scans and 9 prostate scans were performed with PSMA-NBs, 9 tumor scans and 9 prostate scans were conducted using MBs and 3 tumor scans were conducted using Plain-NB. Due to logistical constraints, two imaging sessions utilized PSMA-NBs only (no MBs), as coordinating two injections and an MRI within the same anesthesia session was not feasible. Scans from the first two weeks were excluded from analysis due to the tumors being too small to be adequately visualized, which hindered proper probe positioning.

### Tumor Segmentation and Time-Intensity Curve Analysis

To minimize bias related to contrast enhancement patterns, regions of interest (ROIs) were defined on baseline B-mode images acquired prior to UCA injection, rather than on contrast-enhanced sequences. Because tumor conspicuity and margin delineation on B-mode ultrasound are limited, the baseline tumor region of interest (ROI) was treated as an approximate “tumor-containing” region rather than a definitive histologic boundary; accordingly, we incorporated a tumor-adjacent compartment to explicitly sample the interface between the contoured lesion and surrounding tissue (Figure 2B-C). Two researchers independently delineated the tumor ROI on the baseline B-mode image using pre-specified criteria (hypoechoic and/or heterogeneous regions suspicious for tumor while avoiding clearly identifiable non-tumor structures, such as vessels, when possible). Inter-observer agreement was assessed, and discrepancies were resolved by consensus through joint review to generate a single final tumor ROI per acquisition.

**Figure 2.**
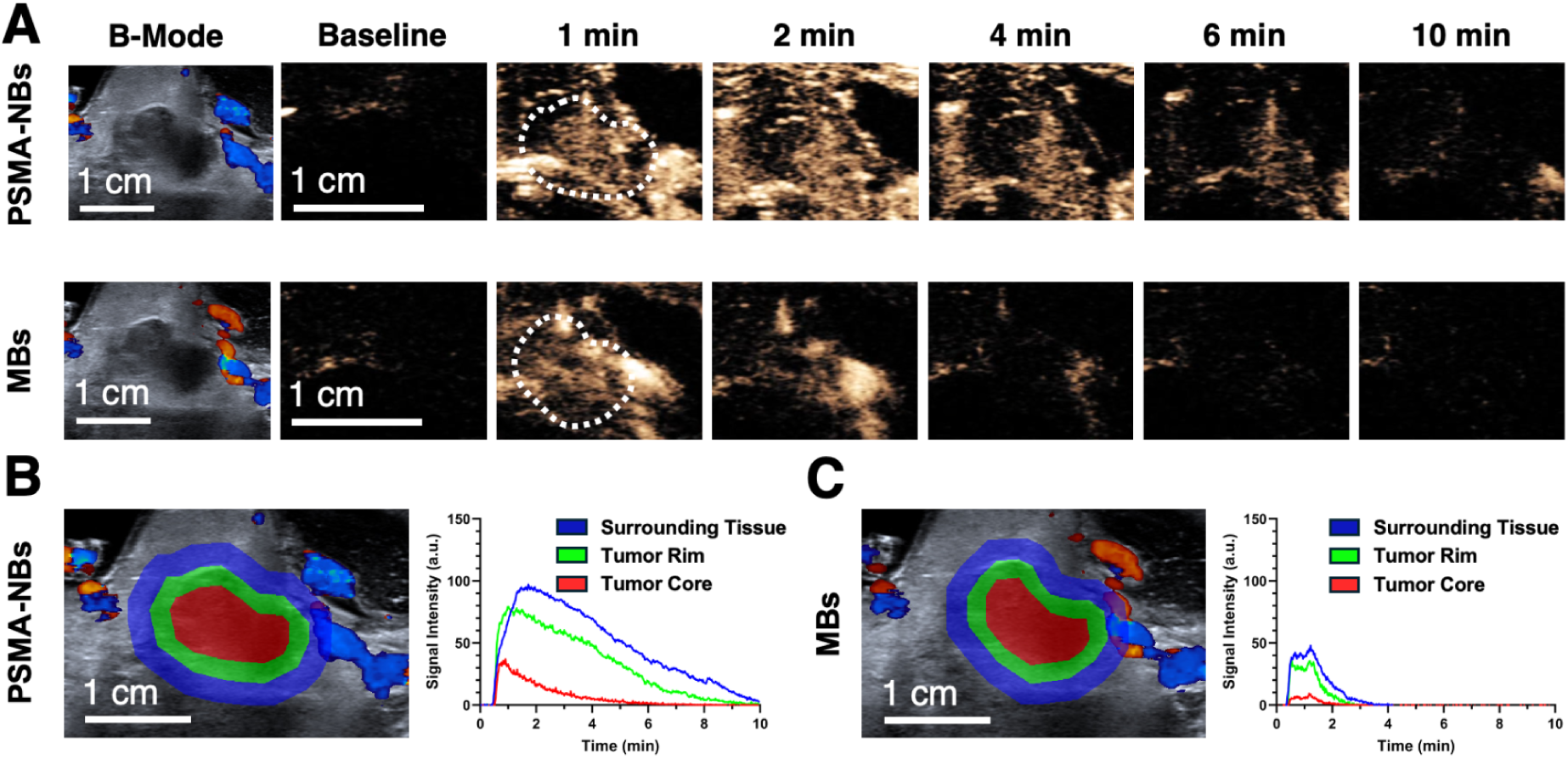
PSMA-NB and MB CEUS output and tumor segmentation strategy. (A) Representative images of B-Mode and non-linear contrast mode *in vivo* of PSMA Nanobubbles (PSMA-NBs) and clinically approved MBs during a 10 min scan in the same tumor of the same animal. (B-C) Example region of interest (ROI) for the tumor core (red), tumor rim (green) and periphery tissues surrounding the tumor (blue) and their respective time-intensity curves (TICs) for (B) PSMA-NBs and (C) MBs.

In order to identify biologically significant spatial heterogeneity while maintaining a reproducible and non-contrast-guided approach, the consensus tumor ROI was subdivided using an automated geometry-based approach, wherein the tumor ROI is subdivided into two equal area compartments consisting of an inner core region representing 50% of the total tumor ROI area and an outer rim region representing the remaining 50%. This approach is consistent with established gradients in perfusion and viability that are typically seen in solid tumors, wherein the outer rim is likely to harbor viable and relatively well-perfused tissue, and the core is likely to harbor nonviable tissue due to lack of vascular supply.^36–40^

Moreover, an ROI for the peritumoral area was created automatically as a band outside the tumor ROI, with a cross-sectional area equal to 20% of the area of the tumor ROI and not overlapping with it. In recognition of the degree of uncertainty in boundary definition based solely on B-mode characteristics, an ROI was created to sample tissue immediately adjacent to the delineated tumor, potentially including a mix of benign prostatic tissue, inflammatory cells, and microscopic tumor.^41,42^

A specific ROI, representing the non-tumor prostate tissue, was manually defined in an area with normal anatomical appearance when possible, avoiding the presence of large vessels and imaging artifacts. All ROIs (tumor core, tumor rim, surrounding/peritumoral tissue, and prostate reference) were then mapped onto the corresponding nonlinear contrast (NLC) cine acquisition obtained during UCA injection within the same imaging session. TICs were generated by calculating the mean NLC signal intensity within each ROI over time, and quantitative time-intensity curve (TIC) parameters were derived from these curves for subsequent comparisons across ROIs and experimental conditions.

For each ROI, the following TIC parameters were calculated: area under the curve (AUC), area under the rising curve (AUC_r_), area under the wash-out curve (AUC_wo_), peak intensity, time to peak (TTP), mean transit time (MTT), and end-frame average signal, as described by Gu *et al.* (Figure 4).^43^ All analysis was carried out using MATLAB^®^ (The MathWorks Inc. Natick, Massachusetts. Version R2025b).

### Size Stratification

To evaluate the influence of tumor size on CEUS parameters, tumors were stratified into two groups based on cross-sectional area: small tumors (≤ 0.55 cm²) and large tumors (> 0.55 cm²). This threshold was derived from biological and anatomical considerations. Specifically, the normal rabbit prostate has an approximate cross-sectional area of 5 cm²; thus, a 0.55 cm² tumor represents roughly 11% of the prostate cross-section. This proportion is consistent with early-stage prostate tumors in humans, which typically occupy approximately 4% of total prostate volume—equivalent to ∼11% of the cross-sectional area assuming a spherical geometry.^44,45^

### Multiparametric Mapping and Decorrelation Time

Multiparametric maps of AUC, decorrelation time, and peak signal intensity were also generated using a pixel-based approach (Figure 8C-D), as described by Wegierak *et al.*^46^ DT is an ultrasound-derived biomarker that captures how long the contrast signal at a given pixel remains similar over time. In practice, higher DT indicates more persistent (slowly changing) contrast signal, consistent with reduced motion and/or prolonged local retention (e.g., extravasation, constrained transport within the tumor interstitium, or agent adherence), whereas lower DT reflects rapidly varying signal, consistent with faster-moving intravascular contrast or freely perfused tissue. DT maps were computed from the NLC image sequence on a pixel-by-pixel basis using the same thresholding approach described by Wegierak *et al.*^46^ (autocorrelation decay to 0.5) generating spatial maps that highlight intratumoral heterogeneity in contrast dynamics beyond intensity-only metrics.

### Histology and Pathologic Correlations

Five weeks post-tumor injection, rabbits were euthanized, and bladder and tumor tissues were harvested. *Ex vivo* imaging was performed using a Maestro *In Vivo* Imaging System (Perkin-Elmer, Waltham, MA, United States) with a blue filter (excitation 445–490 nm, emission filter 515 nm longpass) to detect GFP. Bladder, accessory sex glands, and tumor tissues were fixed in 10% formalin, embedded in paraffin, cut at 5 μm thickness, and stained with hematoxylin-eosin (H&E) following standard protocols. For immunohistochemistry (IHC), tissues were processed by the pathology core, with PSMA expression assessed using a monoclonal antibody against PSMA (ab19071, Ms mAb to PSMA, Abcam, Cambridge, MA, USA).

Slides were digitized using a Nanozoomer S60 scanner (Hamamatsu, Japan). Whole-slide images (WSI) were analyzed in QuPath (v0.5.1) using a Random Trees pixel-classification workflow.^47^ For H&E WSIs, pixel classification was used to segment the tumor into viable tumor, necrosis, and stroma (Figure 6). Using the same QuPath Random Trees pixel-classification approach, PSMA IHC WSIs were segmented into PSMA-positive and PSMA-negative regions to quantify spatial PSMA expression within the tumor (Figure 8). All segmentation outputs were independently reviewed and validated by a veterinary pathologist.

For regional histologic correlations with imaging, each tumor WSI was additionally partitioned into two equal-area compartments based on the histology-defined tumor boundary: an inner core and an outer rim, each comprising 50% of the total tumor area. A “surrounding tissue” compartment was not defined on histology because histopathology was treated as the reference standard for tumor extent and margin definition; therefore, regional analyses were restricted to core–rim compartmentalization within the tumor boundary.

### Statistical analysis

All statistical analyses were performed using Prism^®^ (Version 10.4.1, GraphPad, La Jolla, CA). To compare the average tumor infiltration correlation parameters for PSMA-NBs versus Plain NBs or MBs across the four regions, i.e., surrounding tissue, rim, core, and prostate, we used two-way ANOVA with a mixed effects model. This statistical test was chosen to suitably address the repeated measures, since the same animals contributed data to more than one zone, and to address the missing data points, e.g., when visualization was not possible or when the data were limited to one structure. Geisser–Greenhouse correction was also employed to address the potential violation of sphericity, thereby satisfying the assumptions for the ANOVA test.

For post hoc pairwise comparisons, the Tukey method was used to control family-wise error rates while maintaining sufficient statistical power. When making direct comparisons between PSMA-NBs and MBs within the same tumor zone. Spearman’s or Pearson’s correlation coefficient (r) was calculated to assess the linear relationships between TIC parameters and the percentage of viable tumor tissue.

All values are presented as mean ± standard error of the mean (SEM). Statistical significance was set at *p < 0.05*.

## Results

### PSMA-NB Versus Untargeted NB

To assess the specificity of molecular targeting with PSMA-NBs, a set of studies was performed in which a distinct cohort of animals was imaged with PSMA-NB and Plain-NB during a single imaging session. Since one of the main goals of ligand-based targeting is to improve molecular specificity and prolong the intratumoral retention of the therapeutic agent, the retention-sensitive TIC parameters were given greater emphasis: MTT and AUC_wo_.

When compared to the Plain-NB formulation, the PSMA-NB formulation demonstrated a 21.0% longer MTT value overall across all tumor regions. However, the greatest differences were seen in the rim and surrounding tissue ROI. The wash-out behavior of the formulations was also evaluated by measuring the AUC_wo_. The PSMA-NB formulation was observed to have a greater AUC_wo_ value compared to the Plain-NB formulation. The greatest differences were seen in the rim and surrounding tissue ROI, with 35.0% (SEM: 25.1%) and 25.0% (SEM: 16.0%) greater AUC_wo_, respectively.

**Figure 3.**
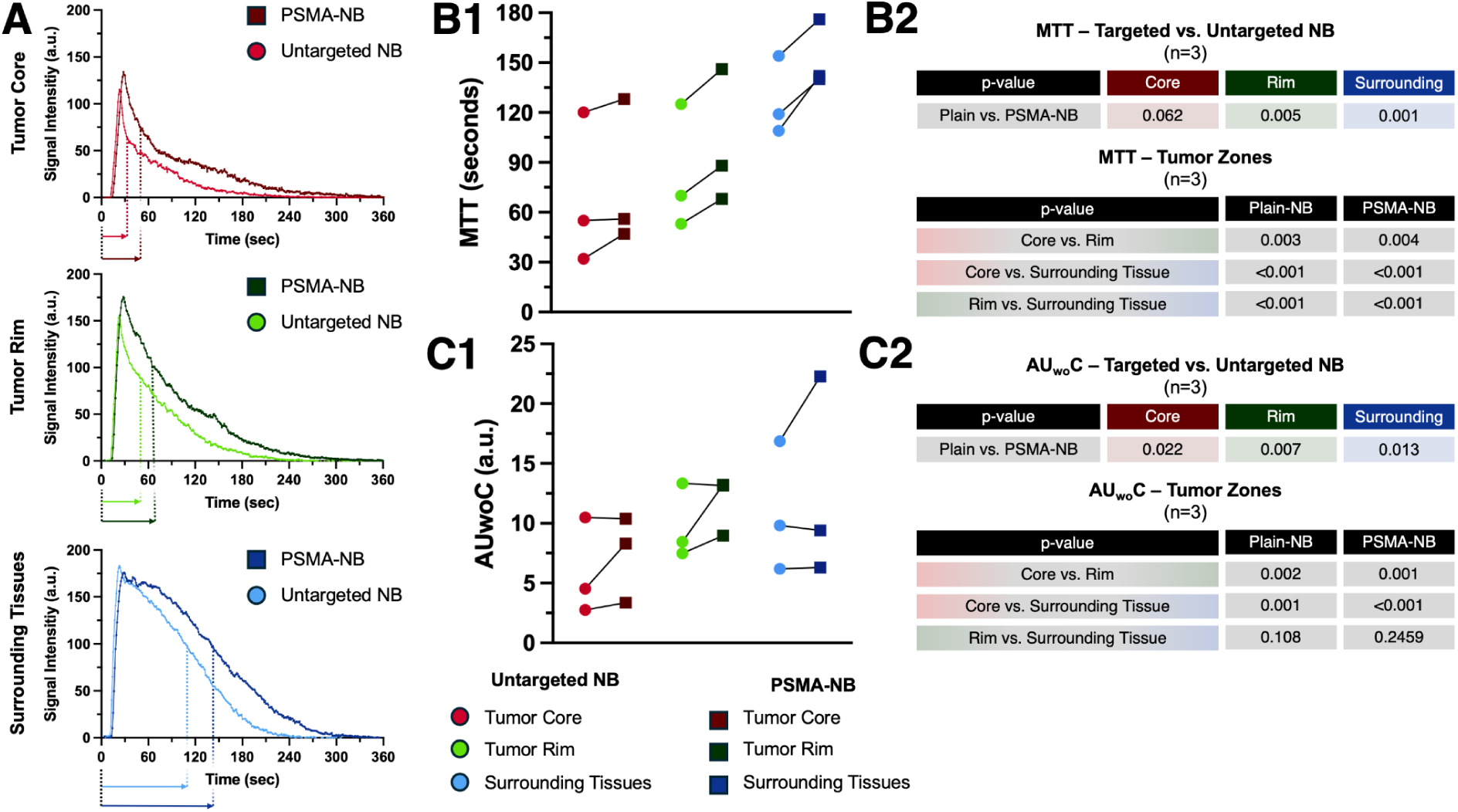
Comparison of PSMA-NB versus Plain-NB (untargeted) kinetics across tumor subregions. (A) Representative TICs from a single rabbit tumor, demonstrating prolonged signal persistence (increased half-life/MTT) of PSMA-NB compared with untargeted nanobubbles (Plain-NB) within the tumor core, rim, and peritumoral (surrounding tissues) ROI. (B) MTT for PSMA-NB versus Plain-NB in each ROI across the three animals imaged with both agents, with corresponding p-values for within-ROI comparisons. (C) AUCwo for PSMA-NB versus Plain-NB across the same ROIs and animals, with corresponding p-values.

### TIC Analyses in Tumor Zones with Comparative Evaluation of PSMA-NBs and MBs

In the PSMA-NB and MB cohort, PSMA-NBs consistently demonstrated higher signal intensity across all tumor zones when compared to MBs. Specifically, PSMA-NBs exhibited a 1.60-fold (p = 0.013) and 1.50-fold (p = 0.016) higher peak signal intensity than MBs in the tumor core and rim, respectively. Additionally, PSMA-NBs showed significantly longer MTTs than those of MBs in both the tumor core (4.20-fold; p = 0.001) and rim (4.50-fold; p < 0.001). A similar effect was observed in the tumor surrounding tissues, where PSMA-NBs displayed a 1.60-fold increase in peak intensity (p = 0.015) and a 5.40-fold increase in MTT (p < 0.001) compared to MBs (Figure 4). Further analysis of TIC parameters revealed that PSMA-NBs exhibited longer MTTs (indicative of extended tissue retention), particularly in the tumor surrounding tissues and rim. In contrast, the tumor core displayed a TIC profile more similar to prostate tissue.

The AUC was significantly higher for PSMA-NBs than for MBs, with the greatest differences observed in the tumor rim (440 ± 80 vs. 60 ± 10 a.u., p = <0.001) and surrounding tissues (580 ± 90 vs. 80 ± 10 a.u., p = <0.001). At the conclusion of the imaging period (end-frame signal), PSMA-NBs remained detectable in the tumor rim (6.0 ± 3.0 a.u.) and surrounding tissues (12.0 ± 5.0 a.u.), whereas no signal from MBs was observed in any tumor zone.

When comparing tumor zones, PSMA-NBs demonstrated differentiation of surrounding tissue from prostate tissue. In the tumor surrounding tissues, PSMA-NBs exhibited a significantly higher AUC (580 ± 90 vs. 250 ± 100 a.u., p = 0.003).. No significant difference in AUC was observed between the tumor rim and the prostate (440 ± 80 vs. 250 ± 100 a.u., p = 0.157) or the tumor core and prostate (250 ± 60 vs. 250 ± 100 a.u., p = 0.999). MBs did not show any significant differences in AUC between surrounding tissues and prostate.

PSMA-NBs also showed significantly higher AUC_wo_ values in the tumor surrounding tissues (470 ± 80 a.u., p = 0.015)) compared to the prostate (210 ± 90 a.u.), whereas the difference between the tumor rim and prostate was not significant (p = 0.255). Similarly, no significant difference was observed in AUCwo between the tumor core and prostate (p = 0.999)). MBs also did not exhibit significant zone-to-prostate differences for AUCwo in the surrounding tissues (53.0 ± 12.0 a.u. vs. 16.0 ± 6.0 a.u., p = 0.999), tumor rim (47.0 ± 11.0 a.u. vs. 16.0 ± 6.0 a.u., p = 0.997) or tumor core (24.0 ± 8.0 a.u. vs. 16.0 ± 6.0 a.u., p = 0.948).

At the end of the imaging period (10 minutes), PSMA-NBs showed higher signal levels in the tumor surrounding tissues compared to the prostate (12.0 ± 5.0 vs. 2.0 ± 1.0 a.u., p = 0.011). However, no significant differences were observed for the tumor rim (7.0 ± 3.0 a.u., p = 0.449) or tumor core (3.0 ± 1.0 a.u., p = 0.999). In contrast, MBs showed no detectable signal in any of the tumor zones, surrounding tissue, or prostate by the end of the 10-minute imaging period.

**Figure 4.**
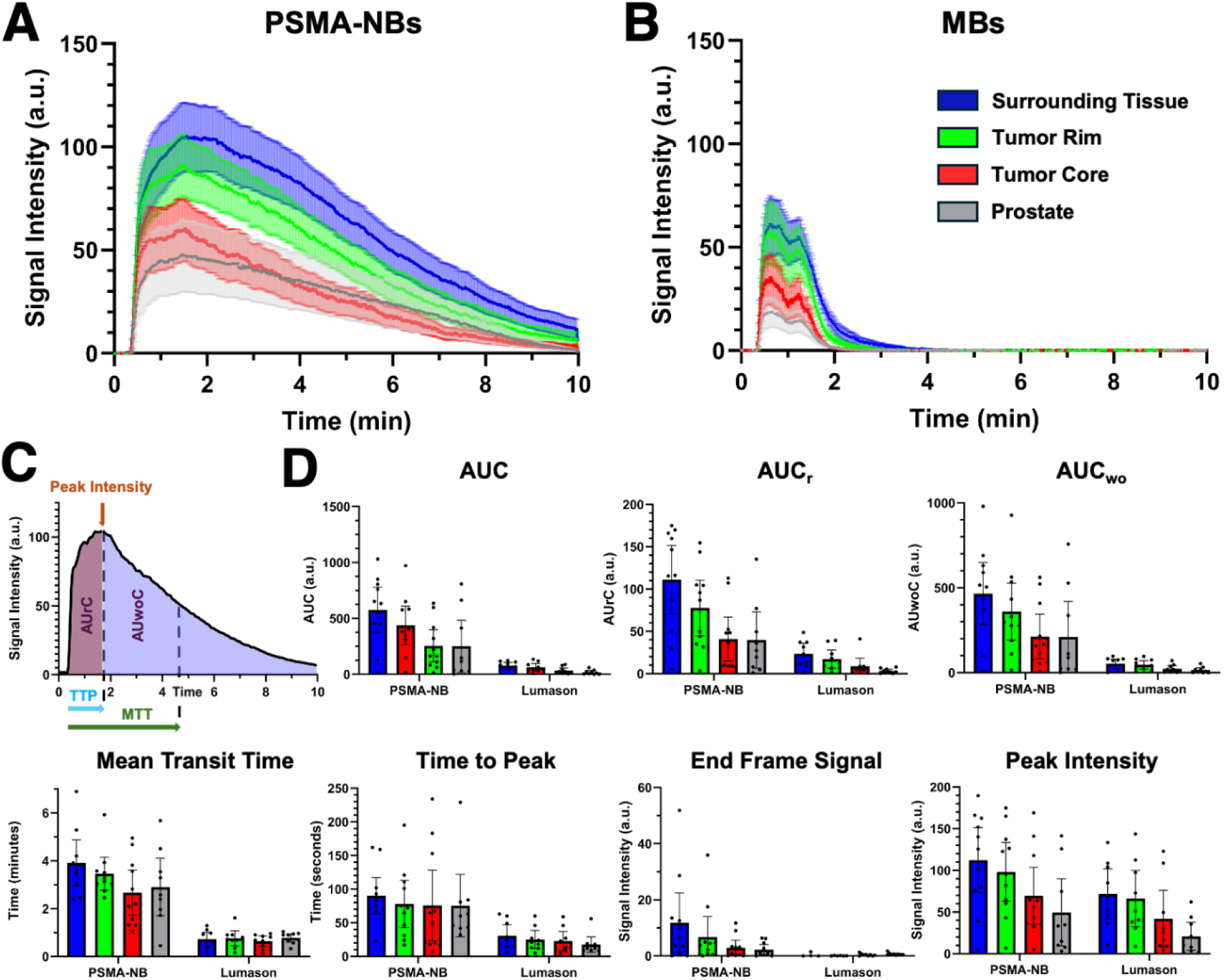
TIC of CEUS in different tumor regions. Time intensity curves (TIC) of the ultrasound non-linear contrast mode average signal in the tumor core (red), tumor rim (green) and periphery tissues surrounding the tumor blue. (A) TIC for PSMA-NBs. (B) TIC for MBs. (C) Graphical representation of TIC parameters and their meaning adapted from Gu *et al.*^43^ (D) Parametric graphs for TIC. AUC: Area under the curve. AUC_r_: Area under the rising curve. AUC_wo_: Area under the wash-out curve. PSMA: Prostate-specific membrane antigen.

### Tumor Size Stratification and Pharmacokinetic Variations

The impact of tumor size on the pharmacokinetics of the UCA was investigated by comparing small tumors defined by an area of ≤0.55 cm² (n = 5) and large tumors defined by an area >0.55 cm² (n = 6). The cut-off was set a priori (Table S1). For all small and large tumors, the PSMA-NBs demonstrated a distinct TIC pattern compared to the MBs, with increased peak signal and retention (longer MTT).

For small tumors, the signal was homogeneous for the different ROI for each type of UCA, with the core, rim, and surrounding tissue ROI for each type of UCA showing similar TIC patterns, and there were no differences between the different regions for the PSMA-NBs or the MBs (Figure 4A1-B1). When the PSMA-NBs were compared to the MBs for the small tumors, the PSMA-NBs maintained the pattern seen for the entire group.

In larger tumors, the kinetics of UCA become non-uniform, indicating the onset of biological zoning. With PSMA-NBs, the TICs in the regions of interest demonstrate a trend where the maximum mean transit time is in the surrounding tissue (4.60 ± 0.60 min), followed by the rim (3.70 ± 0.50 min), and the minimum in the core (2.10 ± 0.40 min) (p < 0.001 for core vs rim and core vs surrounding tissue). With contrast-enhanced MBs, the contrast washes out rapidly in all areas, with some differences in the MTTs in the ROIs, where the rim and core values are similar (0.60 ± 0.20 min and 0.90 ± 0.20 min, p = 0.324), and the values in the surrounding tissue differ from those in the tumor core (p = 0.010). These data indicate that small tumors tend not to demonstrate zonal differences for both contrast agents, while in larger tumors, biological zoning is best demonstrated with PSMA-NBs, maintaining high enhancement in the biologically active areas.

### Correlation of Contrast Enhancement TIC Parameters with Tumor Necrosis

We evaluated radiologic–pathologic correlations between CEUS-derived contrast enhancement metrics and histology-defined tumor viability, stratified by tumor size (Figure 5A and Figure 7A). To ensure temporal alignment between imaging and pathology in this fast-growing model, only the final CEUS acquisitions performed immediately prior to euthanasia (PSMA-NB and MB) were used for correlation analyses.

**Figure 5.**
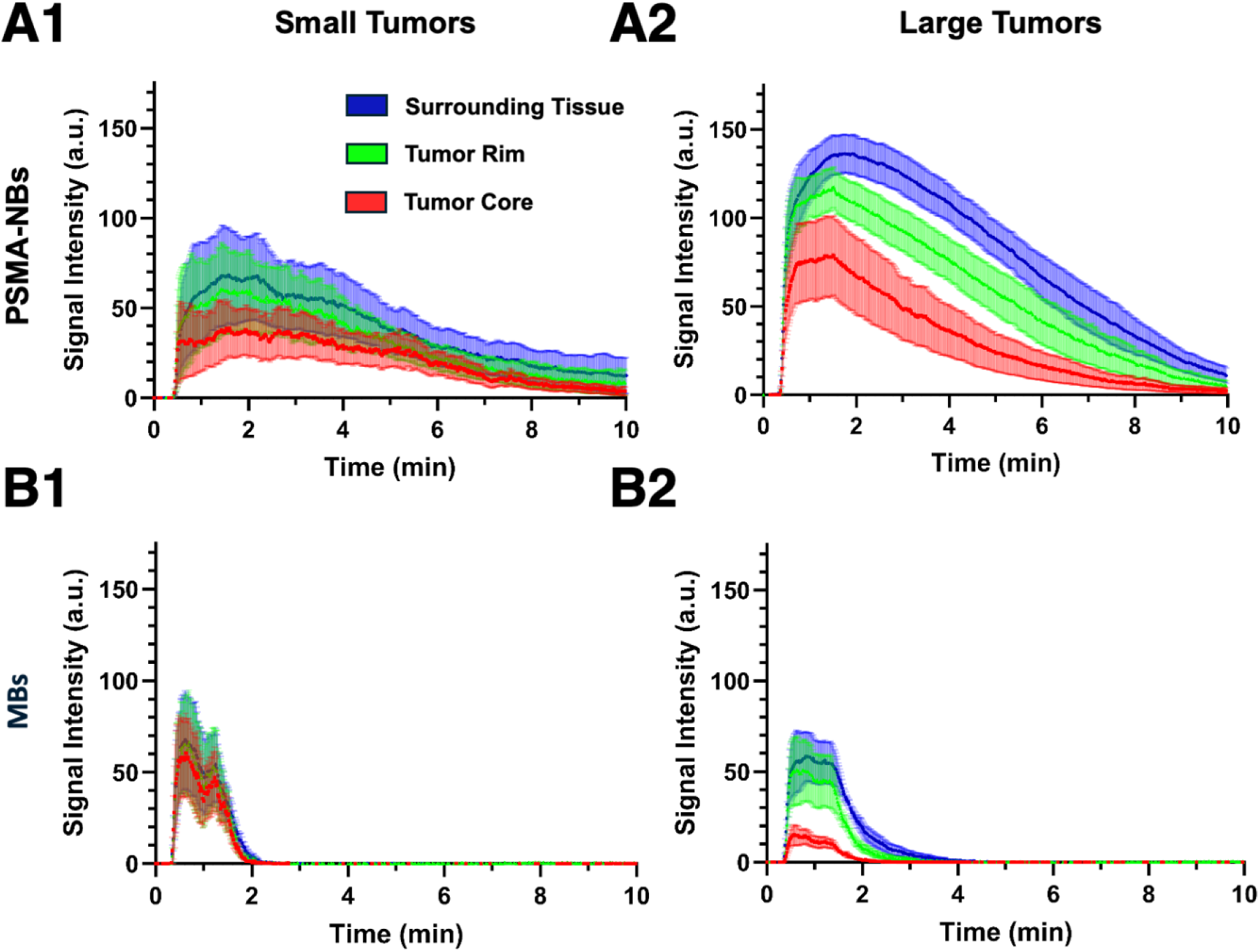
Time-intensity curves (TIC) of the ultrasound non-linear contrast mode average signal in the tumor core (red), tumor rim (green), and periphery tissues surrounding the tumor (blue). TIC for PSMA-NBs in (A1) tumors smaller than 0.55 cm^2^ and (A2) larger than 0.55 cm^2^. (B1) TIC for MBs in tumors smaller than 0.55 cm^2^ and (B2) larger than 0.55 cm^2^.

By the terminal time point, only one animal remained in the small tumor category (≤0.55 cm²), whereas the remaining animals in the PSMA-NB/MB cohort had large tumors (>0.55 cm²). Accordingly, statistical correlations were evaluated for the large-tumor group, where TIC-derived parameters showed size-dependent associations with histologic viability. In large tumors, MTT correlated strongly with viability for both agents, with a stronger association for PSMA-NB (r = 0.96, p = 0.001) than for MB (r = 0.79, p = 0.028). In contrast, AUC_wo_ showed weaker associations with viability in large tumors (PSMA-NB: r = 0.37, p = 0.497; MB: r = 0.16, p = 0.703) (Figure 6).

**Figure 6.**
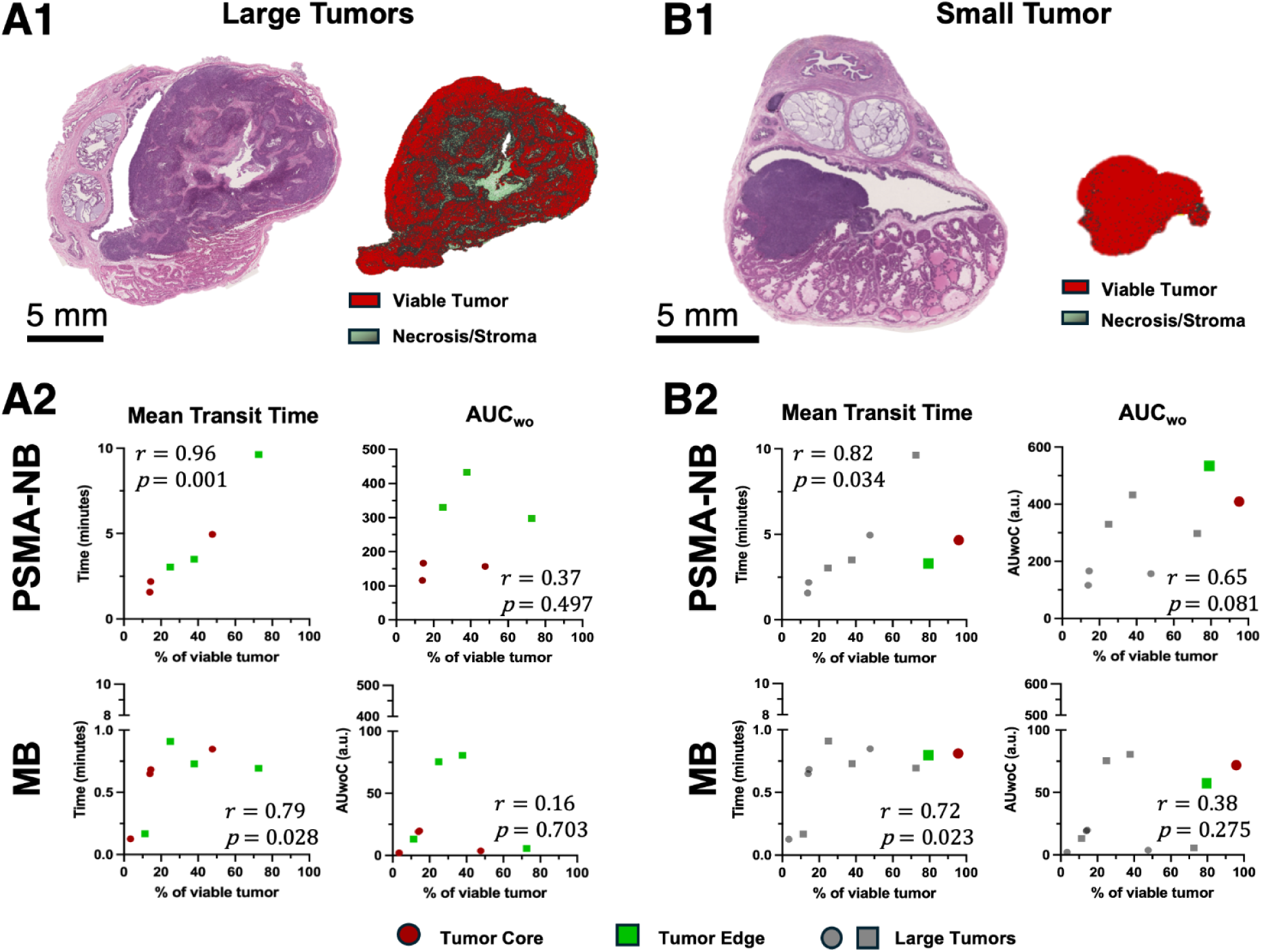
Histopathologic validation of CEUS kinetic metrics across tumor sizes. CEUS–histology correlations were performed using the terminal (euthanasia) imaging session. (A1) Representative whole-section hematoxylin and eosin (H&E) slide from a large intraprostatic tumor (>0.55 cm²), with pixel-based classification into viable tumor, necrosis, and stroma. (A2) Correlation of CEUS-derived kinetic parameters with the percentage of viable tumor quantified from whole-section H&E segmentation in the large-tumor cohort: MTT (left) and AUC_wo_ (right), shown for PSMA-NB (top row) and Lumason® MB (bottom row). Each plot reports the correlation coefficient (r) and p value. (B1) Representative whole-section H&E slide from the single small tumor (≤0.55 cm²), with the same pixel-based classification. (B2) MTT and AUCwo versus % viable tumor for the small tumor (colored symbol) overlaid on the large-tumor cohort (grey symbols) for PSMA-NB (top) and MB (bottom); correlations are computed for the large-tumor cohort, with the small-tumor datapoint shown for reference..

**Figure 7.**
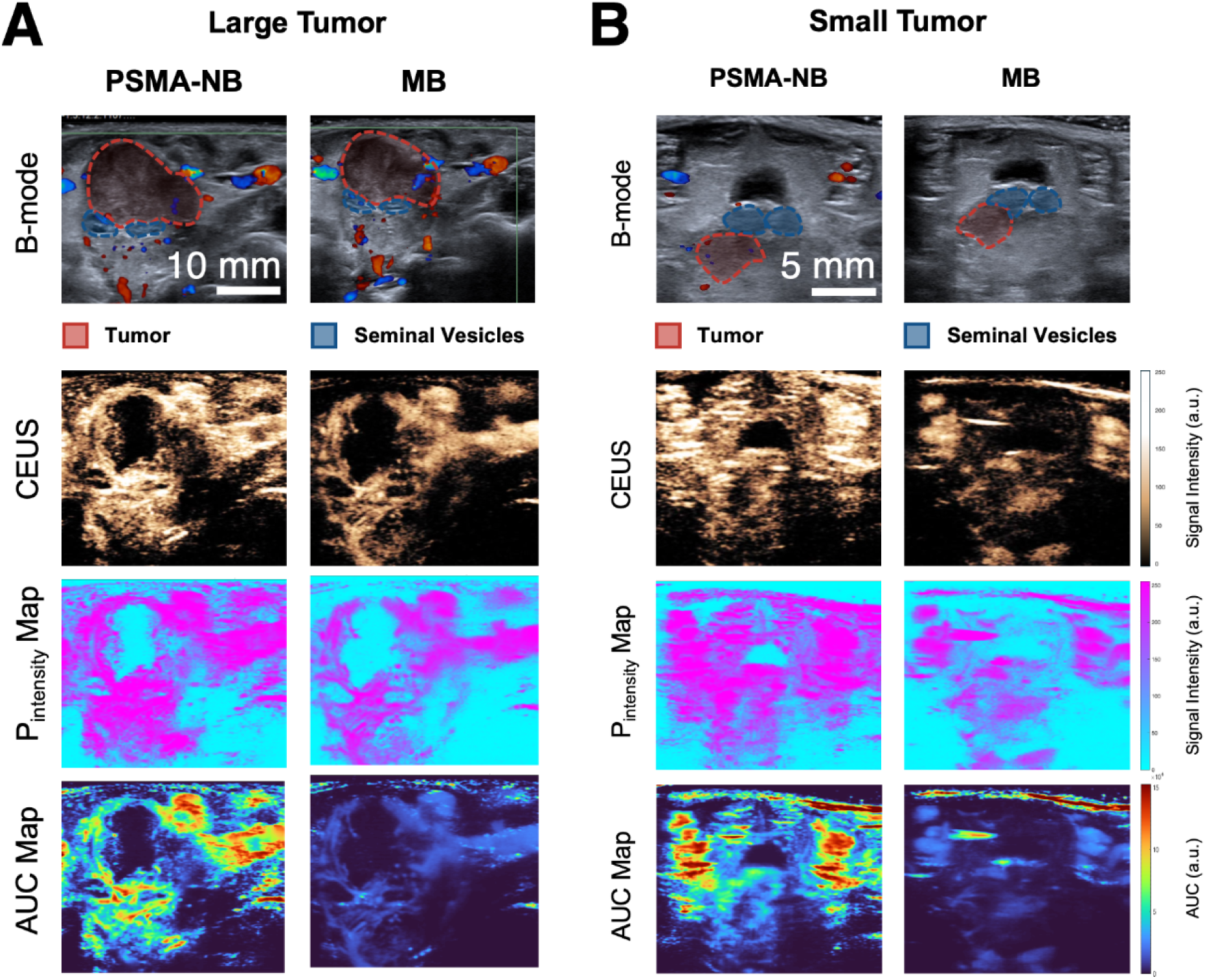
Representative CEUS images and pixel-based parametric maps highlight intratumoral heterogeneity. (A) Large intraprostatic tumor (>0.55 cm²): baseline B-mode anatomy and peak-intensity CEUS frames for PSMA-NB (left) and Lumason® MB (right), with tumor ROI overlay, alongside corresponding pixel-based maps of peak intensity and AUC (AUC) for each agent. (B) Small intraprostatic tumor (≤0.55 cm²): analogous B-mode images, peak-intensity CEUS frames, and peak-intensity/AUC maps for PSMA-NB and MB. Parametric maps illustrate spatially heterogeneous enhancement patterns within and around the ultrasound-defined tumor region.

### Pixel-Based Mapping and Regional Distribution of Contrast Agents

Finally, multiparametric maps were created using a pixel-based approach^46^ (Figure 7C), rather than relying solely on ROI-averaged signal metrics. These pixel-wise maps (peak intensity, AUC, and DT) highlighted spatial heterogeneity in contrast behavior and showed that PSMA-NBs exhibited higher signal intensity within histology-defined viable tumor regions (Figure S3). Notably, DT maps also suggested prolonged signal behavior extending beyond the ultrasound-defined tumor margin.

To relate DT findings to molecular expression, we compared DT values against PSMA positivity quantified from PSMA IHC. In large tumors, rim PSMA% correlated with DT measured in both the ultrasound-defined rim and the surrounding-tissue ROIs, with stronger associations observed when DT was extracted from the surrounding-tissue ROI (Figure 8). This pattern is consistent with boundary uncertainty on ultrasound, whereby histology-confirmed rim tissue may be captured by either ultrasound-defined region. Collectively, pixel-based mapping supported preferential PSMA-NB signal within viable tumor zones and demonstrated that DT-derived spatial patterns tracked PSMA expression at the tumor periphery, including regions adjacent to the ultrasound-defined tumor boundary.

**Figure 8.**
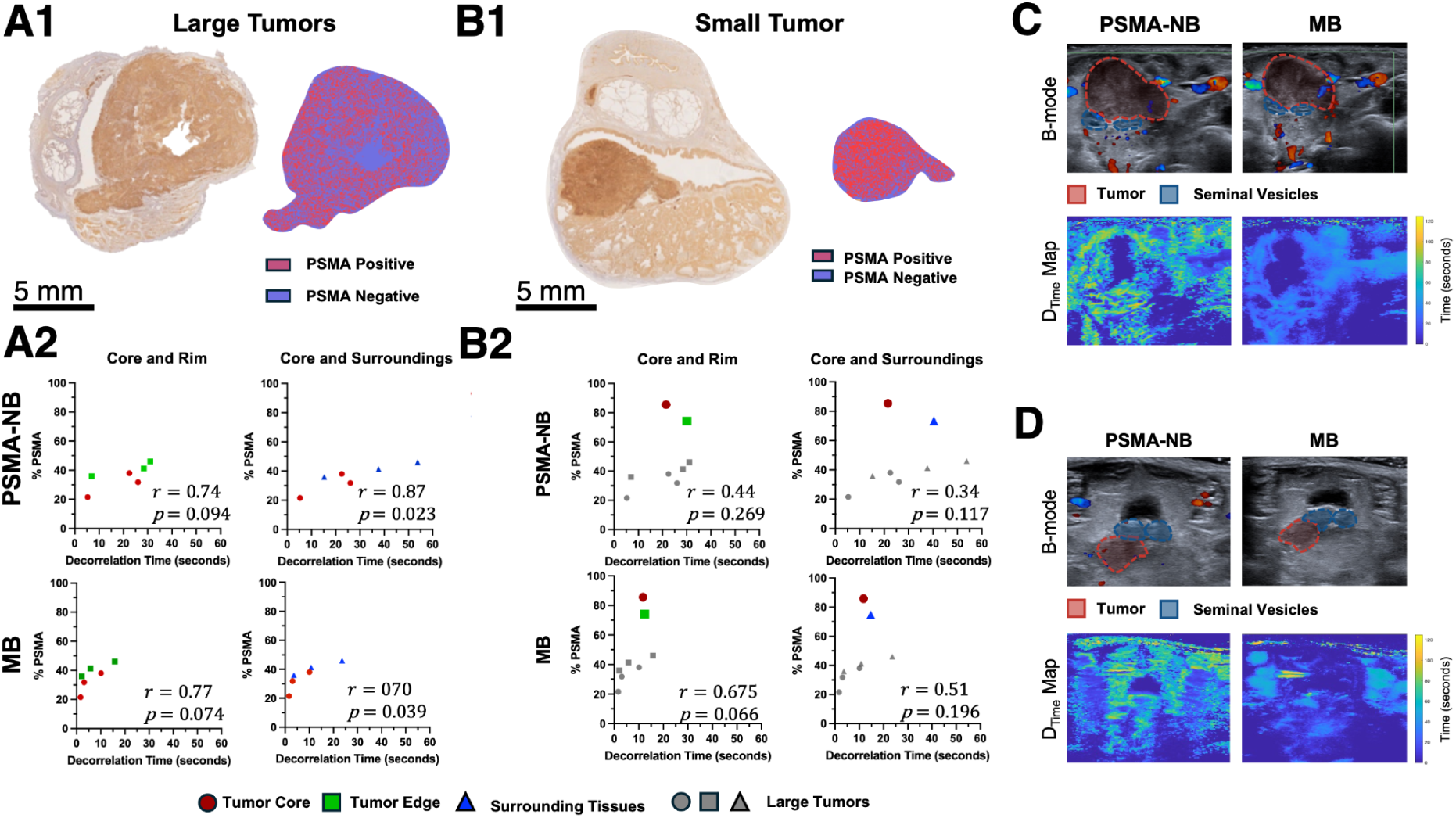
Molecular validation of CEUS decorrelation-time metrics with PSMA immunohistochemistry across tumor sizes. CEUS–IHC correlations were performed using the terminal (euthanasia) imaging session. (A1) Representative whole-section PSMA immunohistochemistry (IHC) from a large tumor (>0.55 cm²), with pixel-based classification into PSMA-positive and PSMA-negative regions (histology segmented into core and rim compartments). (A2) Correlation between percent PSMA positivity and average CEUS decorrelation time (DT) in the large-tumor cohort, shown for PSMA-NB (top row) and Lumason® MB (bottom row). Because PSMA% is available only within the histology-defined tumor boundary (core/rim; ground truth), rim PSMA% is compared against DT measured from (i) the ultrasound-defined rim ROI (left column) and (ii) the ultrasound-defined surrounding-tissue ROI (right column), acknowledging B-mode boundary uncertainty (i.e., true rim tissue may fall within either ultrasound ROI). (B1) Representative PSMA IHC from the single small tumor (≤0.55 cm²), with pixel-based PSMA classification. (B2) DT versus rim PSMA% for the small tumor (colored symbol) overlaid on the large-tumor cohort (grey symbols) for PSMA-NB (top) and MB (bottom). (C) Baseline B-mode images and corresponding DT maps for PSMA-NB and MB from the representative large tumor in (A1). (D) Baseline B-mode images and corresponding DT maps for PSMA-NB and MB from the representative small tumor in (B1).

## Discussion

The results of this study show the capability of PSMA-NBs to differentiate the viable prostate tumor tissues from necrotic tissues and adjacent non-tumoral structures with persistence not possible with conventional, clinically approved MBs. PSMA-NBs also showed improved retention in PSMA-positive tumors relative to untargeted NBs, supporting a measurable contribution of ligand-mediated targeting rather than passive NB behavior alone. PSMA-NBs demonstrated superior signal retention and prolonged presence within the tumor, particularly in the tumor rim and surrounding tissues (e.g. tumor invasion, adjacent fat, seminal vesicles or fibrous tissue), compared to clinically approved MBs and untargeted NB (Video 1). Importantly, these enhanced imaging capabilities were correlated with the percentage of viable tumor tissue, underscoring their potential to enable more accurate diagnosis and targeted therapeutic applications. The enhanced signal is driven by a combination of cell-specific molecular targeting, a high NB concentration per imaging voxel, and strong nonlinear oscillations of PSMA-NBs under insonation, supported by their flexible phospholipid shell a which may enable large volumetric deformation and prolonged acoustic response.^48,49^ Multiparametric imaging further validated that PSMA-NBs selectively accumulate in viable tumor zones, exhibiting prolonged retention and correlation with PSMA expression. These findings support the diagnostic potential of PSMA-NBs in advancing prostate cancer management.

Significantly, in contrast to the earlier studies performed in mice, the present study extends the evaluation of PSMA-NBs to an orthotopic large animal prostate tumor model and includes an untargeted NB control group (Plain-NB). Such a study design will allow for a more rigorous evaluation of whether the PSMA-NB performance will translate to a setting with clinically relevant anatomy, depth of image acquisition, and peritumoral complexity (Figures 3 and 5). Consistent with extensive prior characterization of this formulation—including matched physicochemical properties between targeted and untargeted NBs, validated ligand conjugation, receptor-specific binding in vitro, and improved in vivo tumor retention and accumulation—our rabbit data similarly demonstrate enhanced persistence and retention-sensitive kinetics for PSMA-NBs relative to Plain-NBs, supporting active molecular targeting rather than passive accumulation as the dominant driver of the improved tumor signal.^25,27,28,50^ Although similar results have been observed in murine models, which include orthotopic^28^ and flank models,^25^ these systems have inherent translational constraints due to important differences in prostate histology/anatomy and ultrasound appearance relative to humans. In contrast, the rabbit prostate more closely mirrors human anatomy and ultrasound characteristics (e.g., baseline echogenicity and spatial relationships to adjacent structures),^31^ making it a more relevant intermediate model for assessing tumor conspicuity and peritumoral behavior.

Building on the translational relevance of this orthotopic large-animal model, we observed that PSMA-NBs exhibited significantly longer retention within prostate tissue than clinically approved MBs. This prolonged persistence suggests potential utility beyond tumor conspicuity, particularly for applications that rely on sustained parenchymal enhancement—such as monitoring tissue response after ablative therapies (e.g., high-intensity focused ultrasound [HIFU]), where MBs are currently used for post-procedural assessment.^51^ In this context, the extended prostate signal provided by PSMA-NBs could improve sensitivity for detecting treatment-induced tissue destruction and for tracking post-therapy evolution over time.

Interpretation of enhancement in the “surrounding tissue” regions, however, should be made in the context of a key limitation of anatomic ultrasound: B-mode imaging alone cannot precisely define tumor boundaries, and regions labeled as peritumoral may partially overlap with the biologic tumor rim or include microscopic tumor extension not apparent on grayscale imaging.^41,42,52^ This ambiguity is particularly relevant in prostate cancer, where chronic inflammation in non-tumoral prostate—frequently associated with malignancy—has been linked to increased odds of cancer and is more common in higher-grade disease,^53^ potentially increasing vascular permeability and favoring retention of nanoscale agents. In addition, the rapid tumor growth observed in our model (Figure S4) likely imposed mechanical compression and altered regional perfusion, further influencing contrast-agent distribution and washout.

Consistent with the segmentation and biologic ambiguity of the peritumoral compartment discussed above, we also observed pronounced spatial heterogeneity in contrast enhancement across tumor regions, with signal increasing from the tumor core toward the rim (Video 1). This is consistent with established models of tumor growth in which the rate of growth outstrips the capacity of the vascular supply to meet the metabolic demands of the tumor cells, resulting in central hypoxia, nutrient deprivation, and subsequent areas of necrosis. Thus, there is decreased perfusion-dependent contrast signal in the tumor core compared to the rim.^54,55^ The model also explains the observed phenomenon of less clear delineation of core, rim, and other regions of interest in smaller tumors because they do not develop clear gradients of necrosis or clear viability in the core compared to the rim. Notably, the greatest enhancement is observed in the tissue immediately adjacent to the contoured tumor, which is likely related to peritumoral inflammation and possibly microscopic tumor invasion that is difficult to distinguish using B-mode guidance alone. The association of inflammation/reactive stroma in the peritumoral tissues with aggressive tumor biology and poor outcomes has also been established.^56,57^ Thus, the ability of PSMA-NBs to enhance these tissues could potentially be of value in identifying tumors that have more aggressive invasive potential. These observations support the preferential uptake of PSMA-NBs in the biologically active regions of the tumors and their rims, which is consistent with the potential of these agents to enable characterization of tumors using CEUS.^58^

This is in line with the observed patterns of regional enhancements that have already been described in the above sections. The sustained presence of the PSMA-NBs in the viable tumor tissues as well as in the margin-associated regions is likely to be beneficial in improving the characterization of prostate cancer tissues using molecular CEUS. The sustained presence of the PSMA-NBs in the tissues could also have implications in the context of theranostic approaches that require sufficient residence times to enable the ultrasound-mediated delivery of the therapeutic agents.^59,60^ Notably, the sustained enhancements of the PSMA-NBs in the prostate tissues while concurrently clearing over time can be considered to be in line with the specific targeting of the agents in the tissues. The results of earlier preclinical studies on the PSMA-NB formulations have provided support to the translation of the agents to the human context. These studies have demonstrated favorable stabilities of the agents in physiological media as well as biodistribution patterns that are consistent with the reticuloendothelial system-mediated clearance of the agents. These patterns have included the uptake of the agents by splenic macrophages.^61,62^ Notably, the results of a pilot toxicology study in Sprague-Dawley rats have demonstrated that there were no adverse effects of the agents after the administration of a high dose of 70 mg/kg. The results also revealed that there were mild non-adverse effects on the hematologic parameters as well as spleen tissues that were consistent with the clearance of the agents.^63^

Several limitations need to be addressed while interpreting the results of this study. Firstly, the small sample population (N=9) could be a limitation. Secondly, as a trans-abdominal two-dimensional acquisition was used for imaging, it was not feasible to obtain a single plane from all the time points as the tumors grew. Moreover, as the bowel was not always positioned in a manner that would allow for clear visualization of the tumors during certain time points, variability was also introduced in this aspect. In addition, variability was also introduced in aligning the regions of interest from the imaging modality to the pathology sections for all the tumors. Finally, the rapid progression of the tumor model may not reflect the slower growth kinetics typical of human prostate adenocarcinoma, which could affect translational applicability of necrosis patterns and contrast kinetics.^64^ This decision reflects that “dose” in bubble-based CEUS is not well captured by bubble count alone, particularly when comparing agents with inherently different size distributions. Prior modeling and experimental work indicate that acoustic efficacy scales more closely with total gas volume and size-dependent scattering than with particle number,^65^ and MB concentration–dependent attenuation can further complicate direct “dose equivalence” at higher concentrations. Accordingly, we used volume-matched injections (0.8 mL/kg) to standardize administration conditions and enable a clinically relevant head-to-head comparison across agents. However, because nonlinear response and attenuation effects—especially for MB—can influence TIC-derived parameters, standardized volume matching cannot fully eliminate dosing-related confounding and should be considered a study limitation.^66^ Future studies incorporating gas-volume normalization and/or dose–response designs would further refine quantitative comparisons across formulations.

In conclusion, this orthotopic large-animal study shows that PSMA-NBs preferentially enhance viable tumor and margin-associated regions and demonstrate prolonged retention relative to both clinically approved MBs and untargeted NBs. These findings support continued development of PSMA-NBs as a molecular CEUS approach for prostate cancer characterization and justify future studies using volumetric imaging and more clinically representative disease models to define diagnostic performance and evaluate theranostic applications.

## Data Availability Statement

The data that support the findings of this study are available from the corresponding author upon reasonable request.

## CRediT Author Statement

**Felipe M. Berg:** Methodology, Software, Validation, Formal analysis, Investigation; Data Curation, Writing - Original Draft, Visualization, Project administration; **Eric C. Abenojar:** Methodology, Validation, Formal analysis, Investigation; Data Curation, Writing - Original Draft, Visualization, Project administration; **Pinunta Nittayacharn:** Investigation, Writing - Review & Editing; **Nathan K. Hoggard:** Validation, Formal analysis, Writing - Review & Editing; **Sidhartha Tavri:** Methodology, Investigation, Writing - Review & Editing; **Jing Wang:** Investigation; Data Curation, Writing - Review & Editing; **Felipe Matsunaga:** Methodology, Investigation, Writing - Review & Editing; **Michaela B. Cooley:** Formal analysis, Writing - Review & Editing; **Dana Wegierak:** Software, Writing - Review & Editing; **Xinning Wang:** Methodology, Resources, Writing - Review & Editing; **Thomas J. Rosol:** Validation, Formal analysis, Writing - Review & Editing; **James P. Basilion:** Conceptualization, Methodology, Resources, Writing - Review & Editing, Supervision, Funding acquisition; **Agata A. Exner:** Conceptualization, Methodology, Validation, Formal analysis, Resources, Data Curation, Writing - Original Draft, Visualization, Supervision, Project administration, Funding acquisition

